# Automating UbiFast for High-throughput and Multiplexed Ubiquitin Enrichment

**DOI:** 10.1101/2021.04.28.441860

**Authors:** Keith D. Rivera, Meagan E. Olive, Erik J. Bergstrom, Alissa J. Nelson, Kimberly A. Lee, Shankha Satpathy, Steven A. Carr, Namrata D. Udeshi

**Affiliations:** Broad Institute of MIT and Harvard, Cambridge, MA 02142, USA; Cell Signaling Technology, Danvers, MA 01923, USA

## Abstract

Robust methods for deep-scale enrichment and site-specific identification of ubiquitylation sites is necessary for characterizing the myriad roles of protein ubiquitylation. To this end we previously developed UbiFast, a sensitive method for highly multiplexed ubiquitylation profiling where K-ε-GG peptides are enriched with anti-K-ε-GG antibody and labeled on-antibody with isobaric labeling reagents for sample multiplexing. Here, we present robotic automation of the UbiFast method using a magnetic bead-conjugated K-ε-GG antibody (mK-ε-GG) and a magnetic particle processor. We report the identification of ∼20,000 ubiquitylation sites from a TMT10-plex with 500 μg input per sample processed in ∼2 hours. Automation of the UbiFast method greatly increased the number of identified and quantified ubiquitylation sites, improved reproducibility and significantly reduced processing time. The workflow enables processing of up to 96 samples in a single day making it suitable to study ubiquitylation in large sample sets. Here we demonstrate the applicability of this method to profile small amounts of tissue using breast cancer patient-derived xenograft (PDX) tissue samples.

## Introduction

Ubiquitylation is a highly-conserved protein post-translational modification regulating a wide variety of cellular functions including regulation of protein turnover through the ubiquitin-proteasome system ^1^. The ubiquitylation process is regulated by E1 activating, E2 conjugating, E3 ligating enzymes together with deubiquitinases. Dysregulation of ubiquitylation enzymes and deubiquitinases can lead to aberrant activation or deactivation of pathways involved in many disease processes, notably cancer progression, neurodegeneration and innate and adaptive immune regulation^2^. Drugs targeting the ubiquitin system such as proteasome inhibitors have proven highly successful in the clinic. The development of additional therapeutics targeting this pathway are emerging but remain highly dependent on continued understanding of ubiquitination biology in disease^3–5^.

Liquid chromatography-mass spectrometry (LC-MS/MS) is the leading method for unbiased analysis of protein modifications, including protein ubiquitylation^6,7^. Analysis of ubiquitylated proteins is typically carried out by first using trypsin to generate peptides suitable for LC-MS/MS analysis. Trypsin cleaves proteins at the carboxyl side of lysine (Lys) and arginine (Arg). When ubiquitin is attached to a substrate protein, trypsin digestion leaves a glycine-glycine (GG) remnant on the side chain of Lys residues of tryptic peptides which were formerly ubiquitylated. Antibodies that recognize this di-glycl remnant (K-ε-GG) are used to enrich ubiquitylated (Ub) peptides for analysis by liquid-chromatography-tandem mass spectrometry (LC-MS/MS)^8–11^.

Isobaric chemical tags such as the Tandem Mass Tag (TMT) system are commonly used to compare up to 16 samples within a single experiment and provide precise relative quantitation of peptides and proteins^12–14^. A major limitation of integrating TMT quantitation and ubiquitylation profiling with K-ε-GG antibodies is that these antibodies no longer recognize and enrich peptides when the N-terminus of the di-glycyl remnant is derivatized with TMT. To overcome this limitation, we recently developed the UbiFast method for highly sensitive and multiplexed analysis of ubiquitylation sites from cells or tissue ^15^. The UbiFast method employs an anti-K-ε-GG antibody for enrichment of K-ε-GG peptides followed by on-antibody TMT labeling. Specifically, K-ε-GG peptides are labeled with TMT reagents while still bound to the anti-K-ε-GG antibody which allows the NHS-ester group of the TMT reagent to react with the peptide N-terminal amine group and the ε-amine groups of lysine residues, but not the primary amine of the di-glycyl remnant. In this way, TMT-labeled K-ε-GG peptides from each sample are combined, eluted from the antibody, and analyzed by LC-MS/MS. The UbiFast approach eliminates cell culture restrictions, greatly reducing the amount of input material required when using SILAC-based experiments^8,9,16–22^ and improves Ub-peptide recovery and analysis time compared to in-solution TMT labeling strategies^23^. The sensitivity of the UbiFast method makes it suitable for studies in primary tissue samples.

Although the UbiFast method is highly effective for deep-scale LC-MS/MS analysis of ubiquitylated peptides, the throughput is limited because all workflow steps are manually executed, making the procedure laborious. Additionally, an initial cross-linking step is necessary to covalently couple the antibody to agarose beads, which prevents contamination of antibody in enriched samples ^9^. Further, slight variations during the K-ε-GG peptide enrichment step can result in an increased variability across replicates within a given TMT experiment.

To improve the UbiFast method, here we evaluate and optimize use of a commercially available anti-K-ε-GG antibody supplied irreversibly coupled to magnetic beads (HSmag anti-K-ε-GG). Additionally, we demonstrate that HSmag anti-K-ε-GG antibody enables automation of the UbiFast method on a magnetic bead processor greatly increasing sample processing throughput while maintaining low variation across experimental replicates (Figure 1). We also show that the automated UbiFast method can be easily scaled to process multiple 10-plex TMT experiments in just a few hours. The automated UbiFast method was benchmarked against the previously published, manually executed UbiFast method for profiling patient-derived breast cancer xenograft tissue.

**Figure 1.**
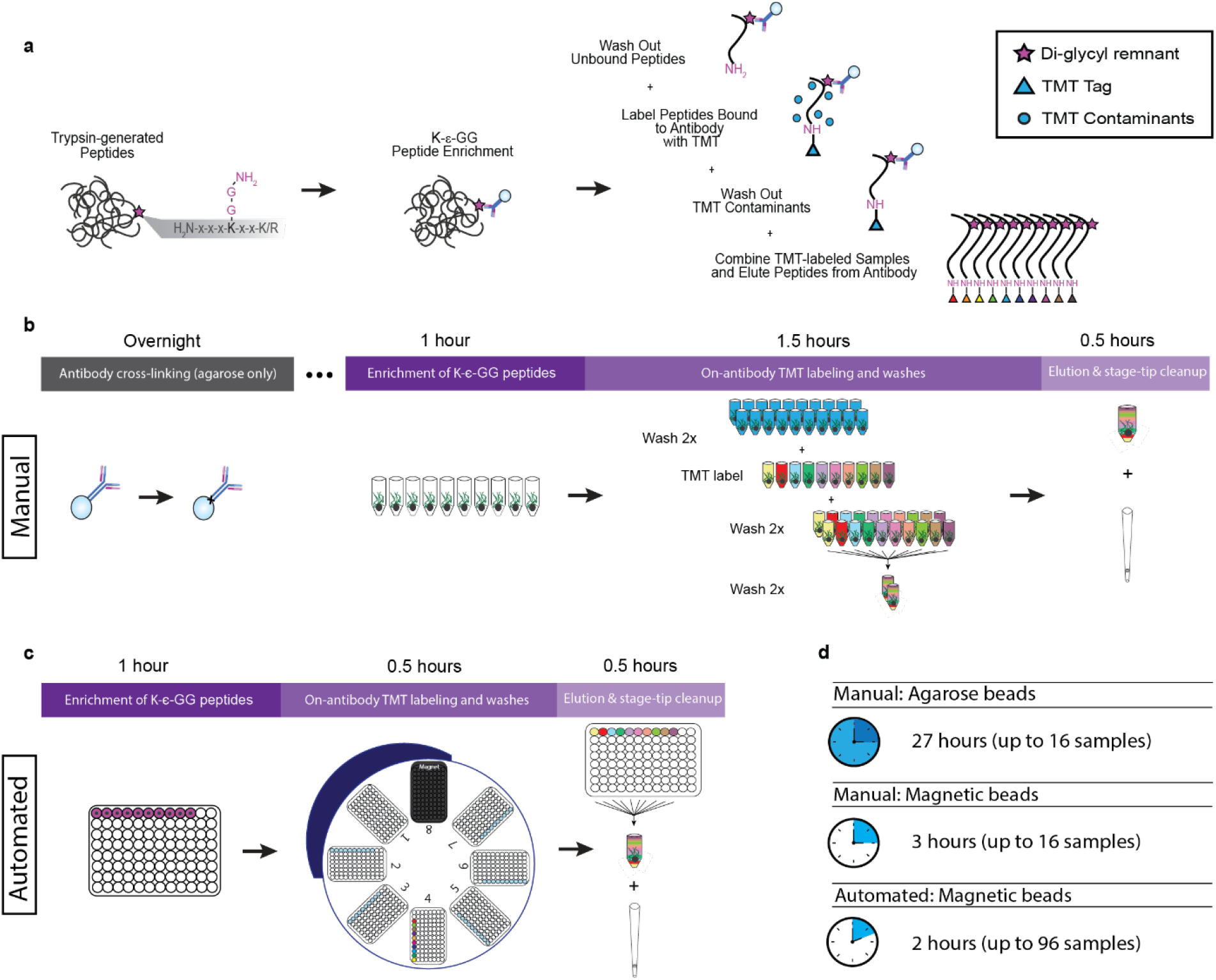
Comparison of methods for on-antibody labeling of K-ε-GG peptides with isobaric reagents. a Workflow of on-antibody labeling of K-ε-GG peptides with isobaric reagents using the UbiFast method. b Manual UbiFast workflow for processing a TMT10 experiment. c Automated workflow of processing samples for a TMT10 experiment using 96-well plates and a KingFisher Flex instrument.

## Experimental Procedures

### In-solution digestion of Jurkat cell lysates

Jurkat cells were grown in suspension with RPMI 1640 medium, glutaMAX supplement (Life Technologies) and 10% heat-inactivated fetal bovine serum (Life Technologies). Cells were pelleted, washed 2x with 1X PBS (pH 7.4), and lysed. Lysis buffer consisted of 8 M urea, 50 mM tris hydrochloride (Tris-HCl) pH 8.0, 150 mM sodium chloride (NaCl) and 1 mM ethylenediaminetetraacetic acid (EDTA). Immediately before lysing the following additives were added to their respective final concentrations: aprotinin (Sigma) to 2 ug/mL, leupeptin (Roche) to 10 ug/mL, phenylmethylsulfonyl fluoride (PMSF) (Sigma) to 1 mM, PR-619 (LifeSensors) to 50 uM and chloroacetamide (CAA) to 1 mM. Cells were lysed for 30 min on ice and then centrifuged for 10 min at 20,000 x g and 4°C. Protein concentration was then determined using a bicinchoninic acid (BCA) assay kit (ThermoFisher Scientific). The lysate was diluted to 8 mg/mL with lysis buffer, dithiothreitol (DTT) was added to a final concentration of 5 mM and it was incubated at room temperature (RT) for 45 min. Next, iodoacetamide (IAA) was added to a final concentration of 10 mM and the lysate was incubated for 30 minutes at RT in the dark. Then the lysate was diluted 1:4 with 50 mM Tris-HCL, pH 8.0. After dilution Lys-C (Wako) was added with an enzyme to substrate ratio of 1:50 and the lysate was incubated for 2 hrs at RT. Trypsin was then added with an enzyme to substrate 1:50 and the lysate was incubated overnight at RT. The following morning neat formic acid (FA) was added to a final concentration of 1% and the sample was centrifuged for 10 min at 10,000 x g to remove urea and undigested proteins. Peptides were cleaned up using a 500 mg Sep-Pak tC18 solid phase extraction cartridge (Waters). Briefly, the cartridge was conditioned with 5 mL of Acetonitrile (ACN) / 0.1% FA and then 50% CAN / 0.1% FA. Equilibration was performed with 4 additions of 5 mL 0.1% trifluoroacetic acid (TFA). Digested peptides were loaded and then washed twice with 5 mL 0.1% TFA. Peptides were eluted with 2 additions of 50% ACN / 0.1% FA. The eluate was frozen at -80°C and dried to completion in vacuo. Peptides were reconstituted in 30% ACN / 0.1% FA and peptide concentration was measured using a BCA assay kit. The peptides were divided into 500 μg aliquots, frozen at -80°C and dried in vacuum. Once dried the peptide aliquots were stored at -80°C until use.

### Label-free comparison of manual immunoprecipitation with agarose vs. magnetic beads

Agarose beads used in all antibody comparisons were from the PTMScan® Ubiquitin Remnant Motif (K-ε-GG) Kit (Cell Signaling Technology, Kit #5562). Preparation of agarose beads included cross-linking as described previously ^15^. All agarose bead washes were performed with immunoprecipitation (IAP) buffer (50 mM MOPS, pH 7.2, 10 mM sodium phosphate, 50 mM NaCl). Magnetic beads for all experiments come from PTMScan® HS Ubiquitin/SUMO Remnant Motif (K-ε-GG) Kit (Cell Signaling Technology, Kit #59322), along with 1X HS IAP Bind Buffer #1 and 1X HS IAP Wash Buffer. Magnetic beads were used as provided, with no additional cross-linking. Input for manual comparisons was 500 μg digested Jurkat peptide, and enrichments were performed in triplicate.

Jurkat samples were reconstituted with 1.5 ml of either IAP buffer for agarose-bead IPs or 1X HS IAP Bind Buffer #1 for magnetic-bead IPs. Reconstituted peptides were confirmed to be pH 7 and were spun at 10,000 x g for 5 min (pH adjusted with 1 M Tris if necessary). After cross-linking, agarose beads were suspended in a slurry of 1:25 beads to IAP buffer from which 62.5 μl slurry (31.25 μg antibody) was used per IP. Magnetic beads are provided in a 20% slurry, and enough was removed for a given experiment to be washed 3x with 1 ml ice-cold 1X PBS, inverted ∼5x with each wash, and resuspended again in original volume. Different magnetic bead amounts were tested for this label-free comparison, including 10 μl slurry (2 μl beads) and 5 μl slurry (1 μl beads) per IP. Beads were aliquoted into clean 1.5 ml snap-cap Eppendorf tubes.

Reconstituted peptides were added to aliquoted antibody beads and incubated at 4 °C for 1 h, gently rotating end-over-end. Buffers were chilled during this enrichment step, and all remaining wash steps were performed on ice when tubes were not being handled. After incubation, agarose beads were centrifuged at 2,000 x g for 1 min and allowed to settle for ∼10 s before removing supernatant as IP flowthrough. Simultaneously, magnetic beads were spun at 2,000 x g for 5 s and allowed to settle for ∼10 s on a magnetic rack before slowly removing flowthrough. Sample flow throughs can be stored at -80 °C for serial analyses.

All washes were performed as quickly as possible. For agarose bead IPs, beads were washed 4x with 1.5 ml PBS, inverting ∼5x with each wash, centrifuging at 2,000 x g for 1 min, letting beads settle and aspirating supernatant. Magnetic beads were washed 4x with 1 ml 1X HS IAP Wash Buffer and 2x with 1 ml HPLC H_2_O, inverting ∼5x with each wash, centrifuging for ∼5 s at 2,000 x g, and allowing beads to be drawn to magnet for ∼10 s before aspirating supernatant. Ubiquitin peptides were eluted from all beads with 50 μl 0.15% TFA for 10 min at RT, gently flicking the beads into solution every 2-3 min. While eluting, 2-punch Empore C18 (3M) StageTips were conditioned and equilibrated 1x with 100 μl methanol (MeOH), 1x with 100 μl 50% ACN/0.1% FA, and 2x with 1% FA.

Upon completion of the first elution, samples were centrifuged quickly at 2,000 x g, placed on a magnetic rack where applicable, and the supernatant containing ubiquitin peptides was loaded onto its designated conditioned StageTip. The elution step was repeated and the supernatant similarly loaded onto the StageTip to be washed 2x with 100 μl 1% FA and eluted with 50 μl 50% ACN/0.1% FA. Desalted eluates were transferred to HPLC vials, frozen at -80 °C, and dried via vacuum centrifugation.

Samples were reconstituted in 9 μl 3% ACN/0.1% FA and 4 μl was analyzed via nanoflow liquid chromatography coupled to tandem mass spectrometry, or LC-MS/MS, using an Easy-nLC 1200 system (Thermo Fisher Scientific) online with a Q-Exactive HF-X mass spectrometer (Thermo Fisher Scientific). Samples were injected and chromatographically separated on a fused silica microcapillary column (360 μm OD X 75 μm ID) with a 10 μm electrospray emitter tip (New Objective) packed to ∼25 cm with ReproSil-Pur 1.9 μm C18-AQ beads (Dr. Maisch GmbH) and heated to 50 °C. Online separation occurred over a 154 min gradient, employing a changing ratio of solvent A (3% ACN/0.1% FA) to solvent B (90% ACN/0.1% FA). Gradient steps as min:% solvent B include 0:2, 2:6, 122:35, 130:60, 133:90, 143:90, 144:50, 154:50, starting at a mobile phase flow rate of 200 nl/min for the first six steps and increasing to 500 nl/min for the final two.

Ion acquisition was performed with a data-dependent analysis method. MS1 scans were collected across a range of 300-1800 m/z at a resolution of 60,000, with an automatic gain control (AGC) target of 3E6 ions and maximum injection time of 10 ms. Within a scan, the top 20 most abundant peaks were picked for higher-energy collisional dissociation (HCD) fragmentation using a collision energy of 28 and isolation window of 0.7 m/z. MS2 spectra were acquired in centroid mode at a resolution of 45,000 with an AGC target of 1E5 and maximum injection time of 150 ms. Peptide matching was set to “preferred”, dynamic exclusion was 20 s, and ions with unassigned charge or charge =1 or >7 were excluded.

### Comparison of TMT labeling efficiency with agarose vs. magnetic beads

A comparison of tandem mass tag (TMT) labeling efficiency of K-ε-GG peptides captured by agarose versus magnetic antibody beads was done in triplicate, using 500 μg Jurkat peptide input. Beads were aliquoted as described above, using 62.5 μl agarose slurry and 5 μl magnetic slurry per IP. Sample reconstitution, incubation with antibody, and flowthrough collection were performed exactly as outlined above. Samples enriched with agarose or magnetic beads were washed once with 1 ml IAP buffer or 1X HS IAP Wash buffer and once with 1 ml 1X PBS or HPLC H_2_O, respectively. With each wash, samples were inverted ∼5 times, centrifuged at 2,000 x g for ∼5 s and allowed to settle on ice or a magnetic rack before aspirating the supernatant.

After washing and immediately before labeling, all beads were resuspended in 200 μl 100 mM HEPES (pH 8.5). Each sample was labeled with a single TMT channel (126C, 127N, 127C for agarose replicates and 128N, 128C, 129N for magnetic replicates), adding 400 μg reagent in 10 μl anhydrous ACN directly to the resuspended beads. Tubes were shaken at 1400 rpm for 10 min at RT. Labeling was then quenched with 8 μl 5% hydroxylamine followed by 5 additional minutes of shaking at 1400 rpm at RT. Excess reagent was washed away 1x with 1.3 ml and 2x with 1.5 ml respective IAP wash buffers, inverting tubes ∼5 times, centrifuging for ∼5 s at 2,000 x g and allowing beads to settle before removing the supernatant. Beads were similarly washed 2x with 1.5 ml 1X PBS (agarose) or HPLC H_2_O (magnetic) before eluting and stage tipping exactly as described above.

Samples were analyzed similarly to the previous label-free experiment, with the reconstitution and liquid chromatography method being identical. Data acquisition was performed using an Orbitrap Exploris 480 mass spectrometer (Thermo Fisher Scientific) with a data dependent analysis method. MS1 parameters were the same as above, with the exception of a 100% normalized AGC target. A precursor fit filter was employed, with a fit threshold of 50% and window of 1.4 m/z. MS2 spectra were collected with an HCD collision energy of 32%, resolution of 15,000, AGC target of 50%, and maximum injection time of 120 ms. Remaining ion acquisition parameters were the same as previously described.

### Developing Enrichment of K-ε-GG peptides using the KingFisher Flex

For all optimization enrichments 500 μg of dried Jurkat peptides were reconstituted in 250 μl PTMScan^®^HS IAP Bind Buffer #1 with a final concentration of 0.01% CHAPS and placed in a sonicator bath for 2 minutes. pH was confirmed to be neutral and then solution was cleared by centrifugation for 5 minutes at 10,000 x g. Magnetic beads from PTMScan® HS Ubiquitin/SUMO Remnant Motif (K-ε-GG) Kit were prepared as described above. After washing beads 3x with ice cold 1x PBS beads were reconstituted to their original volume with HS IAP Bind Buffer #1 and beads were aliquoted into individual wells of a 96 well KingFisher plate 200μl (ThermoFisher Scientific). The solution of cleared peptides was added to the wells containing beads, the plate was covered with aluminum sealing film (Axygen) and rotated end-over-end for 1 hour at 4°C, except where otherwise noted.

In order to determine if CHAPS is a necessary addition to buffers when using the KF we processed triplicate ubiquitin enrichments with all buffers containing either 0.01% CHAPS or all buffers without CHAPS added. To assess the feasibility of performing the K-ε-GG peptide capture step on the KF, enrichments were done in triplicate on the KingFisher for 1 hour at room temperature with medium mixing. Separately, duplicate enrichments were done in 96-well KingFisher plates offline with end-over-end rotating in a cold room at 4°C. The wash buffer used for these experiments was HS IAP Wash Buffer with a final concentration of 0.01% CHAPS.

To test the viability of using lower volumes of beads, 5 uL, 10 μl and 20 μl of slurry were processed in quintuplicate using the label-free KingFisher Flex method described below. Each replicate was injected separately onto the mass spectrometer.

In the interest of assessing the optimal wash buffer, HS IAP Wash Buffer with 0.01% CHAPS was compared with a modified version of this buffer where it was diluted 1:1 with ACN to make 50% ACN, 50% HS IAP Wash Buffer with .005% CHAPS.

For label-free experiments a 6-step KingFisher Flex method was used. Step 1 collects the beads from the incubation plate containing the jurkat peptides and the magnetic beads. For comparisons where incubation of peptides and beads is performed on the KingFisher Flex this step mixes the plate at medium speed for 1 hour. For experiments where the capture of K-ε-GG peptides occurs offline this step mixes the plate at medium speed for 30 seconds and then collects the beads. Steps 2 through 4 are 1 minute washes with 250 μl of washing buffer where the first 15 seconds are on the bottom mix setting and the remaining 45 seconds are set to fast mixing. Step 5 is a wash for 1 minute with 250 μl of HPLC water with medium mixing. The final step is elution in 100 μl of 0.15% TFA for 10 minutes with slow mixing. For all steps a collect count of 5 and collection time of 1 second was used. The eluted peptides were then cleaned up using the stage-tip protocol described above. Similarly, these samples were chromatographically separated and injected onto an Orbitrap Exploris 480 using the same methods described above.

### Comparison of full TMT10 plex experiments manual vs. automated

To evaluate the performance of the automated UbiFast protocol compared to the manual UbiFast protocol using the same PTMScan HS magnetic bead kit we designed three parallel isobaric labeling experiments using TMT10 reagent. For each experiment there were ten process replicates, where each process replicate was 500 μg of dried Jurkat peptides. One of these experiments was performed using the UbiFast protocol^15^. The other two experiments were performed on the optimized automated UbiFast protocol described below. These experiments were identical except one labeled with the TMT10 reagent for 10 minutes and the other labeled with TMT10 reagent for 20 minutes to assess the effect of increased labeling time on labeling efficiency.

For all three experiments the desalted sample was transferred to an HPLC vial, frozen, and dried via vacuum centrifugation. Dried peptides were then reconstituted in 9 μl 3% ACN/0.1% FA and analyzed with back-to-back injections of 4 μl using the same liquid chromatography parameters as above. Ion acquisition was performed with an Orbitrap Exploris 480 (Thermo Fisher Scientific) mass spectrometer in line with a FAIMS Pro™ Interface (Thermo Fisher Scientific). The FAIMS device was operated in standard resolution mode at 100 °C, utilizing the compensation voltages (CVs) of -40,-60, and -80 for the first injection followed by a second injection with CVs of -40, -50, and -70. Remaining parameters were identical to the previous manual TMT-labeling comparison experiment, with the exceptions of utilizing a top 10 method for MS2 spectra collection and an MS2 resolution of 45,000.

### UbiFast protocol using HSmag anti-K-ε-GG reagent

For the manual UbiFast comparison a 10-plex experiment with 500 μg Jurkat peptide input per channel was performed. 5 µL magnetic beads were washed and aliquoted into 10 1.5 ml tubes, and K-ε-GG peptide enrichment was performed as described above. After IP, flow-throughs were collected and saved, all samples were washed with 1 ml 1X HS IAP Wash Buffer followed by 1 ml HPLC H_2_O, inverting ∼5 times before centrifuging for ∼5 s at 2,000 x g and removing the supernatant on a magnetic rack. TMT labeling and quenching was performed identically to the previous single channel experiment, using a full TMT10^™^ 10-plex. Samples were washed first with 1.3 ml followed by 1.5 ml 1X HS IAP Wash Buffer. After washing, each of the 10 TMT-labeled samples were individually resuspended in 90 μl 1X HS IAP Wash Buffer and combined into a single 1.5 ml tube. This combined sample was mixed by inverting ∼5 times and concentrated by spinning in a benchtop centrifuge at 2,000 x g for 5-10 s. Beads were allowed to settle on a magnetic rack and the supernatant was slowly aspirated. The 10 original IP tubes were serially washed with 1.5 ml 1X HS IAP Wash Buffer to collect any remaining beads and this wash was added to the final sample. Two washes with 1.5 ml HPLC water were performed before eluting TMT-labeled K-ε-GG peptides with 150 μl 0.15% TFA for 10 min at RT, tapping the tube every 2-3 min to resuspend the beads. The eluate was collected by spinning the sample briefly, setting the tube in a magnetic rack, and slowly pipetting the supernatant onto a StageTip pre-conditioned as described above. This elution step was repeated once and the sample was StageTip desalted in a manner identical to previous experiments.

### Optimized automated UbiFast protocol using HSmag anti-K-ε-GG reagent

The automated UbiFast experiments were performed as follows: 5 μl of magnetic bead slurry were aliquoted into 10 wells of a 96-well KingFisher plate. 500 μg of dried Jurkat peptides were reconstituted in 250 μl of HS IAP Bind Buffer with 0.01% CHAPS, placed in a bath sonicator for 2 minutes and clarified by centrifugation for 5 minutes at 10,000 x g. The peptide solution was then pipetted into individual wells in a 96-well KF plate containing HSmag anti-K-ε-GG. The plate was covered with aluminum sealing film (Axygen) and rotated end-over-end for 1 hour at 4°C.

The plate containing peptides and HSmag anti-K-ε-GG was then moved to the KF. Immediately prior to beginning the method on the KF, 400 μg of TMT reagent in 10 μl ACN was pipetted into corresponding wells of a 96-well KF plate. Then 190 μl of 100 mM HEPES were added to the TMT containing wells. The KF method for TMT labeling consists of 7 steps (Supplemental Figure 2). Step 1 collects the beads from the incubation plate containing the Jurkat peptides and the magnetic beads after mixing at medium speed for 30 seconds. Step 2 washes the beads with 250 μl of modified HS IAP Wash Buffer [50% ACN / 50% HS IAP Wash Buffer with 0.01% CHAPS] for 1 minute. Step 3 washes the beads with 250 μl of 1X PBS with 0.01% CHAPS for 1 minute. Step 4 is on-antibody TMT labeling with 400 μg of TMT reagent per channel and either 10 or 20 minutes of labeling time. Step 5 is quenching with 250 μl of 2% hydroxylamine for 2 minutes. Step 6 is a final wash with 250 μl of the modified HS IAP Buffer for 1 minute. The final step of the KingFisher Flex protocol is to mix the beads in 100 μl of 1X PBS for 1 minute and then the beads are left in the PBS. For all steps the mixing for the first 15 seconds of the step is set to bottom mix and the remaining time is set to fast. Also, a collect count of 5 and collection time of 1 second was used for all steps. The peptides in PBS were then combined and transferred to a 1.7 mL eppendorf tube and placed on a magnetic rack for 10 seconds. The PBS was removed and 100 μl of 0.15% TFA was added to the beads. This elution was repeated once more and both eluates were loaded onto a stage-tip as described previously.

**Figure 2.**
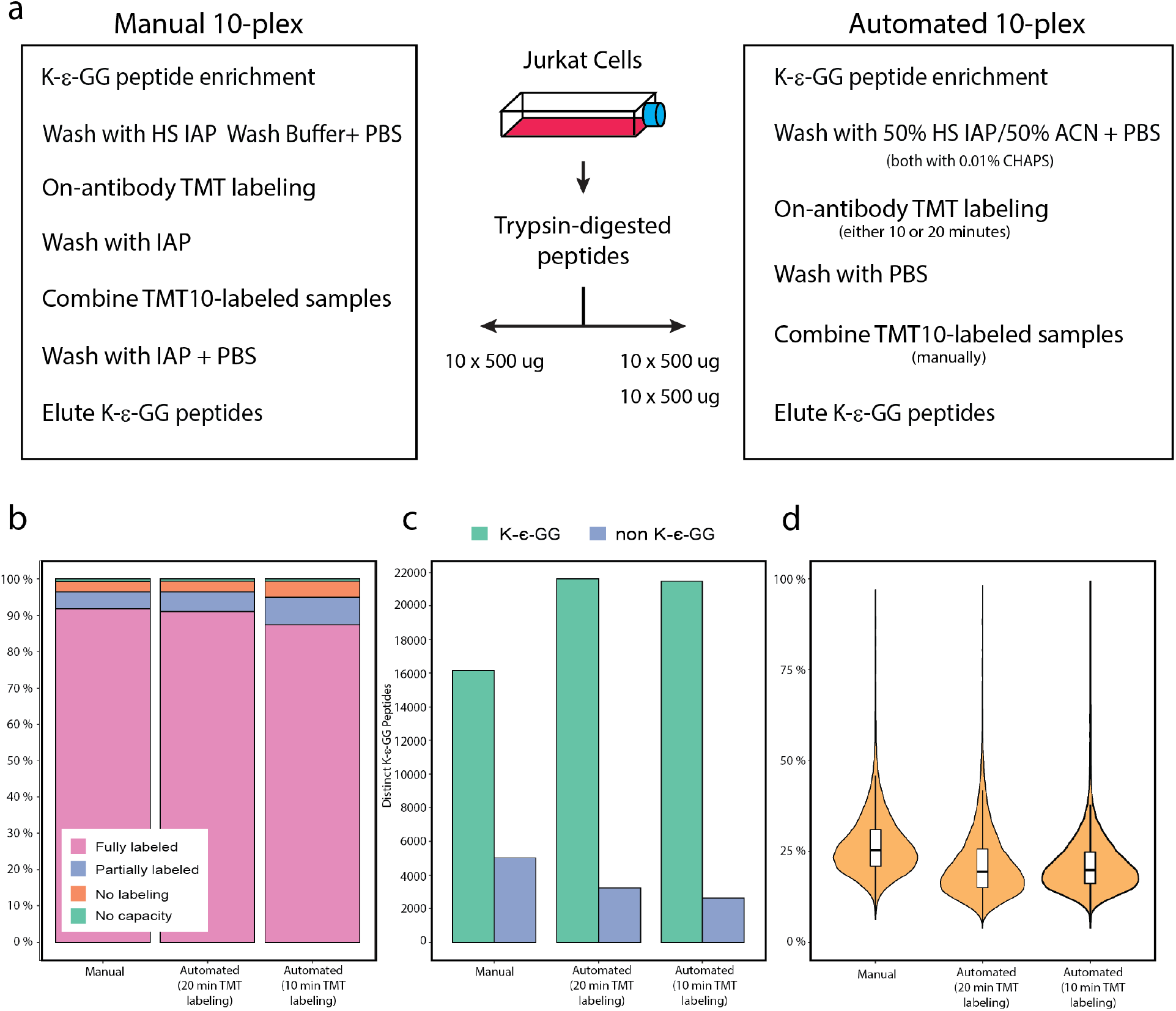
Analysis of K-ε-GG peptide enrichment in a 10-plex format using automation vs. manual workflows. **a** Experimental design of parallel 10-plex ubiquitin enrichment experiments using TMT10 reagents for on-antibody TMT labeling. **b** Stacked bar plots show the percentage of fully, partially and unlabeled K-ε-GG PSM **c** Bar plots of K-ε-GG (green) and non K-ε-GG (blue) distinct peptides identified in each experiment. **d** Violin plots show the distribution of CV’s between differentially labeled TMT channels for each experiment with the inner box and whisker plots showing the median and interquartile range.

### Evaluation of running multiple automated UbiFast experiments simultaneously

To evaluate increased throughput by automatically and simultaneously enriching multiple TMT plexes, 40 replicates of 500 ug dried peptides were enriched for K-ε-GG and labeled as 4 x TMT10 plexes. All steps were carried out in parallel, using the optimized automated UbiFast workflow described above. Briefly, 5 uL of PBS-washed HSmag anti-K-ε-GG bead slurry was aliquoted into 4 × 10 adjacent wells of a 96-well KF plate. Dried 500 ug aliquots of A375 melanoma cell line peptides were each reconstituted in 250 uL of HS IAP Bind Buffer with 0.01% CHAPS, vortexed and shook at 1200 rpm for 5 min, and clarified by centrifugation for 5 minutes at 10,000 g. The peptide solutions were then pipetted into each well containing the magnetic bead slurry. The plate was covered with aluminum film and rotated end-over-end for 1 hour at 4°C. After incubation the plate was processed on the KF where washing, on-antibody TMT labeling and collection of beads in PBS occured. Each set of K-ε-GG enriched peptides were combined by TMT10 plex into 1.7 mL Eppendorf tubes and placed in a magnetic rack for 10 seconds. The PBS was removed and 100 uL 0.15% TFA was added to the beads to elute. The elution was repeated once and both eluates were loaded onto stage-tips as previously described. Desalted samples were analyzed via FAIMS-LC-MS/MS as previously described.

### Processing and analysis of comparative reference tissue

Processing of WHIM2 and WHIM16 patient-derived xenografts (PDX) for ubiquitylome analysis was described previously ^15^. Briefly, frozen tissue from each model was lysed, digested and aliquoted into 500 μg aliquots. Five replicates of WHIM2 and 5 replicates of WHIM16 were included in a single TMT10-plex experiment. The digested tissue was processed using the optimized automated UbiFast protocol and LC/MS analysis was performed exactly as detailed in the previous section.

### Data Analysis

Mass spectrometry data was processed using Spectrum Mill (Rev BI.07.04.210, Agilent Technologies). For all samples, extraction of raw files retained spectra within a precursor mass range of 600-6000 Da and a minimum MS1 signal-to-noise ratio of 25. MS1 spectra within a retention time range of +/- 45 s, or within a precursor m/z tolerance of +/- 1.4 m/z were merged. MS/MS searching was performed against either a human RefSeq database for Jurkat samples or a human and mouse RefSeq database for analysis of PDX samples. Digestion parameters were set to “trypsin allow P” with an allowance of 4 missed cleavages. The MS/MS search included fixed modification of carbamidomethylation on cysteine. For TMT quantitation experiments TMT10 was searched using the partial-mix function. Variable modifications were acetylation of the protein N-terminus, oxidation of methionine and remnant GG on lysine. Restrictions for matching included a minimum matched peak intensity of 40% and a precursor and product mass tolerance of +/- 20 ppm. Peptide matches were validated using a maximum false discovery rate (FDR) threshold of 1.2%. TMT10 reporter ion intensities were corrected for isotopic impurities in the Spectrum Mill protein/peptide summary module using the afRICA correction method which implements determinant calculations according to Cramer’s Rule^31^.

### Experimental Design and Statistical Rationale

For the statistical analysis of PDX models, each protein ID was associated with a log2-transformed expression ratio for every sample condition over the median of all sample conditions. After median normalization, a 2-sample moderated T test was performed on the data to compare treatment groups using an internal R-Shiny package based in the limma library. P-values associated with every protein were adjusted using the Benjamini-Hochberg FDR approach^32^.

## Results

### Comparison of magnetic- and agarose-bead antibody reagents for manual enrichment of K-ε-GG peptides

The PTMScan® HS Ubiquitin/SUMO Remnant Motif (K-ε-GG) Kit (#59322) contains the same monoclonal antibody as the original PTMScan Ubiquitin Remnant (K-ε-GG Kit) (#5562), with the key difference that the new kit contains magnetic beads instead of agarose and bind and wash buffers have been optimized to maximize sensitivity and specificity of K-ε-GG peptide enrichment. The antibody is coupled to the beads using chemistry that does not affect the epitope binding regions. Since this HSmag anti-K-ε-GG formulation does not require an initial chemical cross-linking step to covalently couple the antibody to affinity beads, 1-2 days of procedural time are saved^9,23,24^. The magnetic bead formulation also provides the means to transfer ubiquitin enrichment protocols to a magnetic particle processor for automating and increasing the throughput of sample handling steps.

To characterize the performance of HSmag anti-K-ε-GG, the enrichment of K-ε-GG peptides was directly compared to enrichment with agarose-bead anti-K-ε-GG antibody in a label-free manner by LC-MS/MS. Enrichment of K-ε-GG peptides was completed in triplicate from 500 μg of tryptic peptides derived from Jurkat cells using 5 µl and 10 µl of HSmag anti-K-ε-GG slurry vs. 8 µl of agarose-bead anti-K-ε-GG antibody and enriched peptides were analyzed by LC-MS/MS (Supplemental Figure 1 and Supplemental Data 1). Previous work has shown 8 µl anti-K-ε-GG agarose-bead reagent to be optimal for enrichment of K-ε-GG peptides^25^. An average of 8662 K-ε-GG PSMs were identified with 5 μl of HSmag anti-K-ε-GG, 9284 K-ε-GG PSMs were identified with 10 μl of HSmag anti-K-ε-GG and 6643 K-ε-GG PSMs were identified with 8 μl anti-K-ε-GG agarose-bead antibody. Using 5 μl or 10 μl of HSmag anti-K-ε-GG improved recovery of K-ε-GG PSMs compared to 8 μl anti-K-ε-GG agarose-bead antibody by 30% and 39%, respectively. The relative yield, or the percentage of K-ε-GG PSMs identified relative to the total number of PSMs identified in the sample, was 51% with 5 μl of HSmag anti-K-ε-GG, 57% with 10 μl of HSmag anti-K-ε-GG and 32% with 8 μl anti-K-ε-GG agarose-bead antibody. These results show that HSmag anti-K-ε-GG beads outperform anti-K-ε-GG agarose-bead antibody reagent for the enrichment of K-ε-GG peptides using as little as 5 μl of magnetic beads. Although 10ul of HS mag anti-K-ε-GG beads minimized variability, it provided only a relatively small increase in the number of identified K-ε-GG PSMs vs. the use of 5 μl of HSmag anti-K-ε-GG slurry, indicating that the lower amount of beads is efficient and cost effective.

### Evaluation of on-antibody TMT labeling using HSmag anti-K-ε-GG

The UbiFast method utilizes on-antibody TMT labeling for isobaric labeling experiments ^15^. To evaluate the performance of on-antibody TMT labeling using HSmag anti-K-ε-GG antibody, Jurkat peptides (500 μg in triplicate) were enriched using HSmag anti-K-ε-GG or agarose-bead anti-K-ε-GG antibody and labeled with TMT reagent as previously described ^15^ (Supplemental Data 2). For this evaluation, a single channel of TMT reagent was used to label each replicate. We identified an average of 8008 and 5289 K-ε-GG PSMs with 5 µL magnetic and 8µL agarose-bead antibody reagent, respectively (Supplemental Fig 1b).

To analyze the efficiency of TMT labeling we evaluated the number of K-ε-GG PSMs fully TMT labeled, partially TMT labeled and unlabeled. A peptide is considered fully labeled if the N-terminal amine group (-NH_2_) and the ε-amino group on lysines (if present) have reacted with the TMT reagent. Partially labeled PSMs have one amine group that has reacted with TMT and at least one amine group that has not reacted with the TMT reagent. For the agarose bead experiments, 91% of the K-ε-GG PSMs were fully labeled and 5% were partially labeled. For the magnetic bead experiments, 91% of the K-ε-GG PSMs were fully labeled and 5% were partially labeled (Supplemental Fig 1c). The overlap of K-ε-GG sites identified using HSmag anti-K-ε-GG and anti-K-ε-GG antibody was >52% (Supplemental Figure 1d). Each method also identified a subset of distinct Ub sites. These results demonstrate that on-antibody TMT labeling works effectively with HSmag anti-K-ε-GG and HSmag anti-K-ε-GG outperforms anti-K-ε-GG antibody for identifying K-ε-GG peptides following enrichment and on-antibody TMT labeling.

### Optimization of automated workflow for the UbiFast method

To increase the throughput and reproducibility of the UbiFast method, we sought to reduce the number of manual sample processing steps in the method by automating the HSmag anti-K-ε-GG washing and on-antibody TMT labeling steps using the KingFisher Flex (KF), a magnetic bead processor capable of processing 96-well plates. The KF operates by using magnetic rods to collect magnetic beads from a 96-well plate and transfer them into another 96-well plate. Following offline K-ε-GG peptide capture in a 96 well plate, the plate was loaded onto the KF where antibody beads were washed and K-ε-GG peptides labeled with TMT reagent while still on HSmag anti-K-ε-GG beads (Supplemental Figure 2).

First, we evaluated how efficiently HSmag anti-K-ε-GG transfer from one 96-well plate to a second 96-well plate on the KF. Previous work showed that adding low concentrations of CHAPS to buffers prevents magnetic beads from sticking to the plates and being lost during KF processing steps ^26^. We compared automated UbiFast with and without addition of 0.01% CHAPS to wash buffers and found that CHAPS significantly improves the automated protocol, identifying 1.9-fold more K-ε-GG peptides compared to no CHAPS addition (3329 vs 1752 K-ε-GG PSMs) (Supplemental Figure 3a, Supplemental Data 3). Omitting CHAPS from wash buffers clearly resulted in residual HSmag anti-K-ε-GG beads being left in the plates after the completion of the KF run. These experiments confirmed that CHAPS improves the movement of magnetic beads in this protocol, increasing the number of K-ε-GG PSMs by 89% and decreasing variability due to loss of beads and should be added to all buffers used on the KF platform.

**Figure 3.**
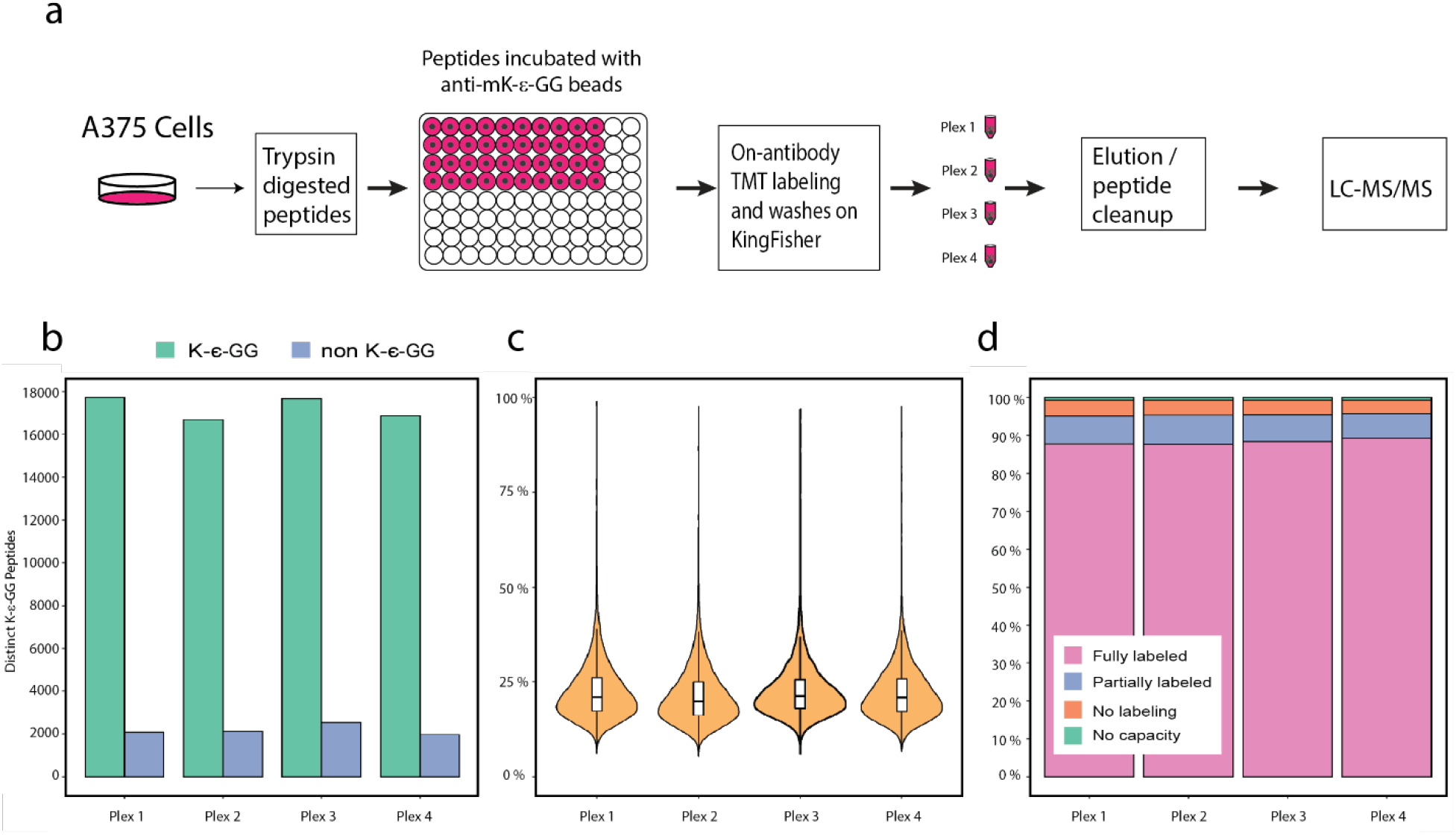
Parallel processing of 4 x TMT10 experiments using the automated UbiFast method. a Schematic of experimental workflow for processing multiple experiments in parallel. b Stacked bar plots show the percentage of fully, partially and unlabeled K-ε-GG PSMs. c Bar plots of K-ε-GG (green) and non K-ε-GG (blue) distinct peptides identified in each experiment. d Violin plots show the distribution of CV’s between differentially labeled TMT channels for each experiment with the inner box and whisker plots showing the median and interquartile range

We next tested if capture of K-ε-GG peptides with HSmag anti-K-ε-GG could be completed on the KF. In the manual protocol, the enrichment step is complete at 4°Cwith end-over-end rotation^9^. The KF does not have a cooling unit and cannot rotate plates end-over-end. Therefore we sought to confirm whether the enrichment step should be performed offline as is done in the manual protocol or on the KF at RT with mixing. To evaluate the capture step, two 96-well KF plates were prepared, each with three wells containing 500 μg Jurkat peptides and the HSmag anti-K-ε-GG. One 96-well plate was rotated end-over-end in a cold room at 4°C and the other plate was incubated on the KF at room temperature with mixing set to ‘medium’ (Supplemental Figure 3b, Supplemental Data 4). A large drop in PSMs was observed when the incubation was performed at RT on the KingFisher, identifying 3308 K-ε-GG PSMs compared to 6215 K-ε-GG PSMs with end-over-end rotation at 4°C. These results indicate that offline end-over-end rotation at 4°C is preferable for binding of K-ε-GG peptides to the HSmag anti-K-ε-GG. Performing this step offline will not add any additional time or labor since the samples and beads are already in the 96-well plate, however, future studies could explore placing the KingFisher in a cold room at 4°C.

Next we determined the optimal HSmag anti-K-ε-GG amount for K-ε-GG peptide enrichment on the KF. Previous work showed that decreasing the amount of agarose-bead anti-K-ε-GG antibody increased the relative yield and overall recovery of K-ε-GG peptides^9^. Given the previous findings, we titrated the amount of HSmag anti-K-ε-GG to determine which bead volume would produce the highest overall number and the highest relative yield of K-ε-GG peptides. We performed quintuplicate enrichments using 5 μL, 10 μL and 20 μL of HSmag anti-K-ε-GG and compared the overall number of K-ε-GG PSMs and relative yield (Supplemental Figure 3c, Supplemental Data 5). The number of identified K-ε-GG PSMs was 6519, 5261 and 4540 for 5, 10 and 20 μL of HSmag anti-K-ε-GG, respectively. The relative yield of K-ε-GG PSMs was 29%, 23% and 20% for 5, 10 and 20 μL of HSmag anti-K-ε-GG, respectively. Enrichment with 5 μL HSmag anti-K-ε-GG provided the best overall number of K-ε-GG PSMs and relative yield of K-ε-GG PSMs.

We noted that the relative yield of K-ε-GG PSMs to total PSMs was much lower for HSmag anti-K-ε-GG processing on the KF relative to manual processing (29% vs 51%). We hypothesized that the decrease in relative yield of K-ε-GG peptides could be due to insufficient bead washing because the volume of each wash step was reduced from 1 mL in the manual protocol to 250 µL on the KF to accommodate the maximum volume of the 96-well plate. To reduce the number of non-K-ε-GG peptides resulting from automated processing, 50% ACN was added to the wash buffer to increase stringency. To evaluate the utility of ACN addition to the wash buffer, we performed a label-free experiment where K-ε-GG peptides were enriched from 500 μg Jurkat peptides where bead washing was completed with or without 50% ACN added to the HS IAP Wash Buffer (Supplemental Figure 3d, Supplemental Data 6). Addition of 50% ACN to the washing buffer increased both the relative yield of K-ε-GG peptides and the absolute number of identified K-ε-GG peptides. An average of 7210 K-ε-GG PSMs with 57% relative yield and 5694 K-ε-GG PSMs with 29% relative yield were identified with and without 50% ACN added to the wash buffer, respectively. Taken together, we find that use of 5 μL of HSmag anti-K-ε-GG bead slurry and inclusion of 0.01% CHAPS and 50% ACN in wash buffers provides the best performance of the UbiFast method on the KF.

### Manual vs. automated TMT 10-plex experiment

The Ubifast method is designed to be used with multiplexed samples^15^. To evaluate the performance of the automated UbiFast workflow, we carried out a head-to-head comparison using HSmag anti-K-ε-GG where both automated and manual UbiFast methods were used to analyze 10 process replicates of peptides from Jurkat cells, with 500 ug peptide input per replicate using a different TMT10 reagent for each replicate. Due to the additional time it takes to prepare and aliquot TMT reagents in a 96-well plate for multiplexed experiments, we performed two independent automated TMT10 UbiFast experiments comparing 10 minutes and 20 minutes on-antibody TMT labeling time to determine if a longer TMT labeling time is needed due to reagent hydrolysis during preparation steps (Figure 2a). For manual and automated experiments, we assessed the efficiency of TMT labeling, relative yield of K-ε-GG peptides and the variability between differentially labeled TMT channels (Figure 2b, 2c, 2d, Supplemental Data 7).

Using the manual UbiFast protocol, we identified 16,141 distinct K-ε-GG peptides with 97% of peptides being labeled with TMT and a relative yield of 75%. The automated UbiFast experiment identified 21,468 with 88% relative yield and 21,601 distinct K-ε-GG peptides with 87% relative yield using 10 and 20 minutes of on-antibody TMT labeling time, respectively. Increasing on-antibody TMT labeling time from 10 min to 20 min increased the TMT labeling efficiency from 95% to 97%. The reproducibility of automated UbiFast experiments was higher than manual UbiFast, with median coefficients of variation across process replicates of 20%, 19%, 25% for automated UbiFast 10min, automated UbiFast 20 min, manual UbiFast, respectively. These data indicate the automated platform washes enriched K-ε-GG peptides and performs on-antibody TMT labeling with high reproducibility, eliminating the need for manual processing.

### Performing multiple experiments in parallel using the automated UbiFast method

Automating the UbiFast method reduces the hands-on time for processing a single multiplexed TMT experiment, but the greatest impact would be in the ability to scale the number of multiplexed TMT experiments that could be processed simultaneously. To evaluate the effects of processing multiple TMT experiments, we designed an experiment in which 4 x TMT10 experiments were processed in parallel (Figure 3a). Briefly, a batch of peptides digested with trypsin from A375 cells were aliquoted into 40 × 500 µg aliquots. The 40 aliquots were split into 4 groups of 10 and processed on the KF as described above with each group of 10 aliquots processed as separate TMT10 experiments. To assess performance across all 4 experiments, we evaluated the number of distinct K-ε-GG peptides, the coefficient of variation for each experiment and the labeling efficiency (Figure 3b, 3c, 3d, Supplemental Data 8). We identified 17,718, 16,704, 17,668 and 16,862 distinct K-ε-GG peptides in experiments 1 through 4, respectively. The median coefficient of variation for each plex was 20.8, 19.8, 21.2 and 20.8 and the percentage of fully labeled K-ε-GG PSMs was 95%-96% for all 4 plexes. These results indicate that the automated UbiFast protocol enables multiple multiplexed TMT experiments to be processed simultaneously with high reproducibility, saving up to 2 hours of laboratory time for each additional TMT experiment.

### Application of automated UbiFast with HSmag anti-K-ε-GG to analysis of tumor samples

Tumor tissues of two breast cancer patient-derived xenograft (PDX) models, basal subtype (WHIM2) and luminal subtype (WHIM16) have been extensively characterized in the proteogenomic space, having been shown to harbor subtype-specific signatures^15,27,28^. To evaluate our automated UbiFast method for profiling ubiquitylomes of small amounts of tumor tissue, we applied the workflow to enrich ubiquitylated peptides from five replicates each of WHIM2 and WHIM16 using 0.5 mg of peptide per sample in a TMT10-plex experiment (Figure 4a). The correlation of replicates within the automated UbiFast dataset employing HSmag anti-K-ε-GG beads was high, with median Pearson correlations of 0.76 and 0.73 for basal and luminal subtypes, respectively (Figure 4b, Supplemental Data 9). In total, 14,211 distinct K-ε-GG peptides were identified and quantified using this method. Ubiquitylation data was compared to previously published results acquired on the sample PDX models, but employing UbiFast in a manual mode using agarose K-ε-GG antibody to evaluate the overlap of K-ε-GG sites and conservation of canonical biological differences between basal and luminal breast cancer models. Among these identified K-ε-GG sites, we see a high degree of overlap at the site level (61%) between datasets (Figure 4c). To assess whether biological changes were conserved across UbiFast workflows, we performed gene set enrichment analysis (GSEA) on both datasets using the GO Biological Process Molecular Signature Database^29,30^. Gene Sets which were significantly changing in both datasets are shown in Figure 4d with the top 5 gene sets enriched in the luminal and basal subtypes from the Udeshi et al. 2020 study highlighted. It is clear that the gene sets which were significantly enriched with the automated method show the same trends as those enriched by Udeshi et al. 2020.

**Figure 4.**
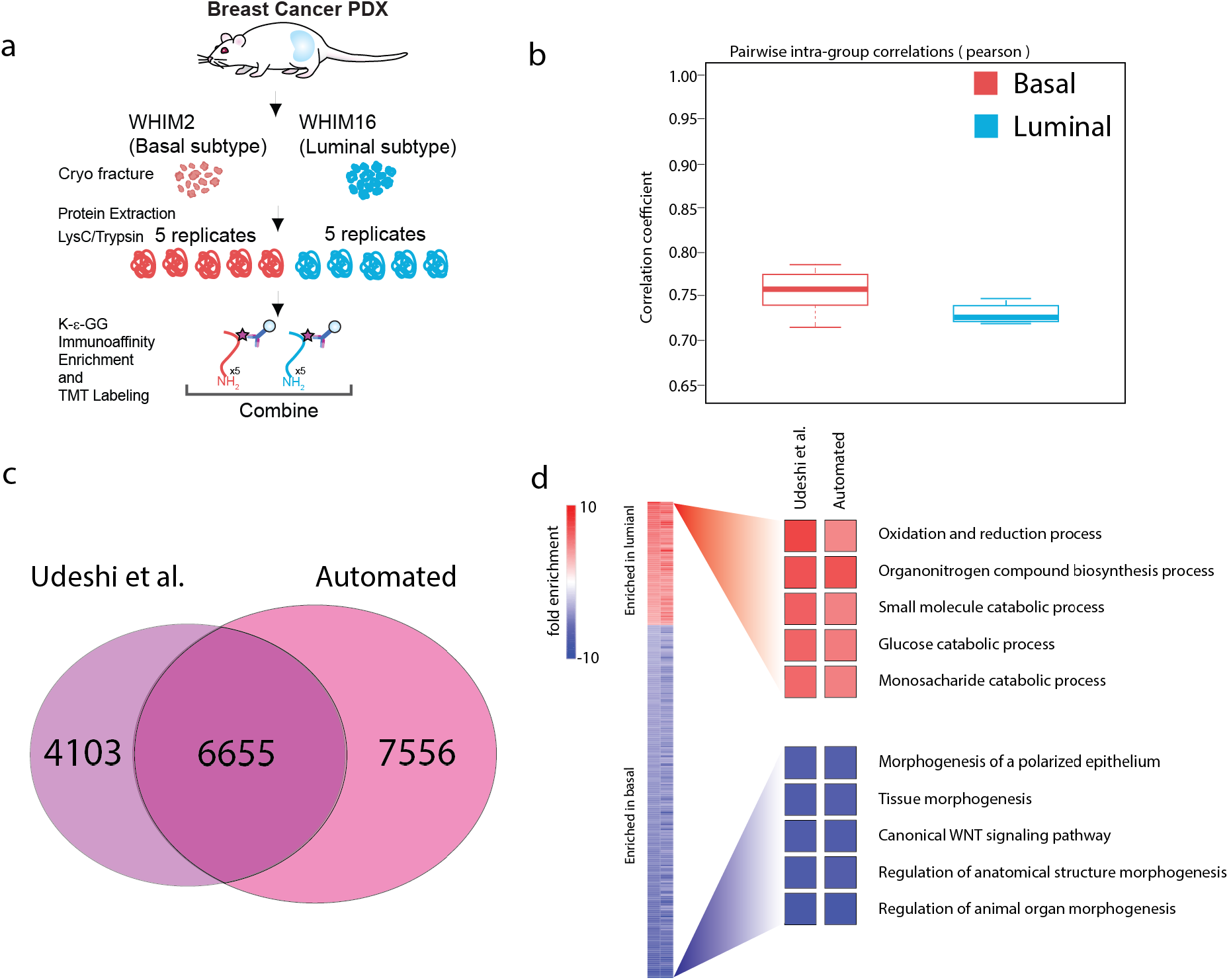
Comparison of the ubiquitylomes identified from our automated workflow vs manually workflow ^15^. a Schematic diagram showing experimental design and process used to enrich K-ε-GG modified peptides from the Luminal and Basal breast cancer PDX models. b Venn Diagram showing overlap of K-ε-GG sites identified in each dataset. c Box of whisker plots of Pearson correlation between replicates of Basal and Luminal PDX models for each dataset. d Heatmap of all genesets present in both the Udeshi et al dataset (left) and the current, automated dataset (right) with the top 5 most enriched gene sets in the Udeshi et al dataset highlighted.

## Discussion

Here we present an automated UbiFast workflow using magnetic-bead K-ε-GG antibody and a robotic magnetic particle processor with a 96 well plate format to increase sample analysis throughput of the UbiFast method while decreasing labor required and chances for human error (Figure 1). Removing the need for a highly trained person to perform these enrichments should make ubiquitylation site profiling accessible to more laboratories. Using the manual implementation of the UbiFast method, it takes more than 1 day of processing time to prepare 10 samples (1 × 10-plex experiment) for injection on the mass spectrometer because the antibody must first be cross-linked to agarose beads followed by quality control^28^. Magnetic HSmag anti-K-ε-GG supplied in the PTMScan® HS Ubiquitin/SUMO Remnant Motif (K-ε-GG) kit are provided with the antibody already linked to the magnetic beads and do not require additional cross-linking, simplifying the method and reducing the total processing time to 3 hours per 1 × 10-plex experiment. Importantly, the magnetic HSmag anti-K-ε-GG reagent identified more Ub-peptides with greater reproducibility than the agarose anti-K-ε-GG reagent in TMT experiments. Automation of the UbiFast method on the KF reduces sample processing time per plex from 3 hours to 2 hours and allows for parallel processing of up to 9 x TMT10-plex experiments. Future efforts will be aimed at extending to higher-plex isobaric labeling methods such as TMTPro (Thermo Fisher), enabling 6 × 16-plex experiments to be processed in parallel. The KF platform may be used to automate the enrichment of other PTMs, such as phosphorylated tyrosine and acetylated lysines as magnetic bead reagents become available in the future.

## Acknowledgements

This work was supported by the National Cancer Institute (NCI) grants U24CA210986, U01CA214125, and U24CA210979 to S.A.C., Swiss National Science Foundation (SNF) Sinergia grant CRSII5_186405 to S.A.C., Dr. Miriam and Sheldon G. Adelson Medical Research Foundation to S.A.C. and N.D.U. and by a SPARC Award to N.D.U. from the Broad Institute of MIT & Harvard (#800373)

## Authors contribution

K.D.R., M.E.O., S.A.C. and N.D.U. conceived the study; K.D.R., M.E.O., and E.J.B. performed experiments. K.D.R., M.E.O., E.J.B., A.J.N., K.A.L., S.S., S.A.C. and N.D.U. contributed to experimental design, data analysis, and data interpretation. K.D.R., M.E.O., S.A.C. and N.D.U. generated figures and wrote the manuscript with input from all authors.

### Data availability

Raw mass spectrometry data will be made publicly available in MassIVE upon acceptance of the manuscript. This article contains supplemental data.

---

**Supplemental Figure 1.**
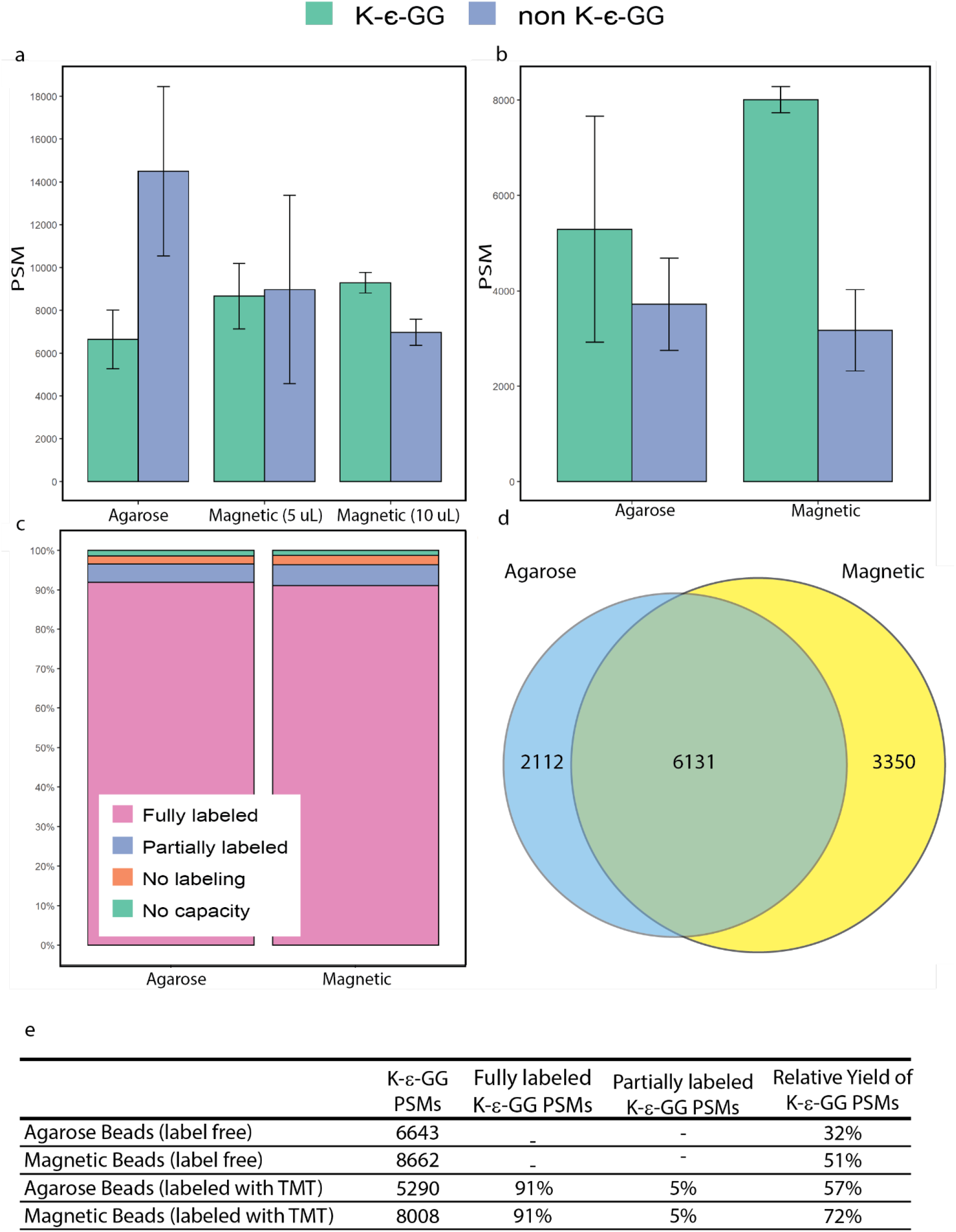
Comparison of K-ε-GG peptide enrichments with antibody coupled to either agarose or magnetic beads. a Bar plots show identification of K-ε-GG and non K-ε-GG PSMs for label free enrichments with agarose and magnetic beads (n=3). b Bar plots show the identification of of K-ε-GG and non K-ε-GG PSMs from enrichments with on-antibody labeling with single channel TMT (n=3). c Stacked bar plots show the percentage of fully labeled, partially labeled, unlabeled and sequences with no capacity for labeling. d Venn diagram showing overlap of peptides identified with agarose and magnetic beads e Table showing all raw values, including enrichment specificity for previously described experiments.

**Supplemental Figure 2.**
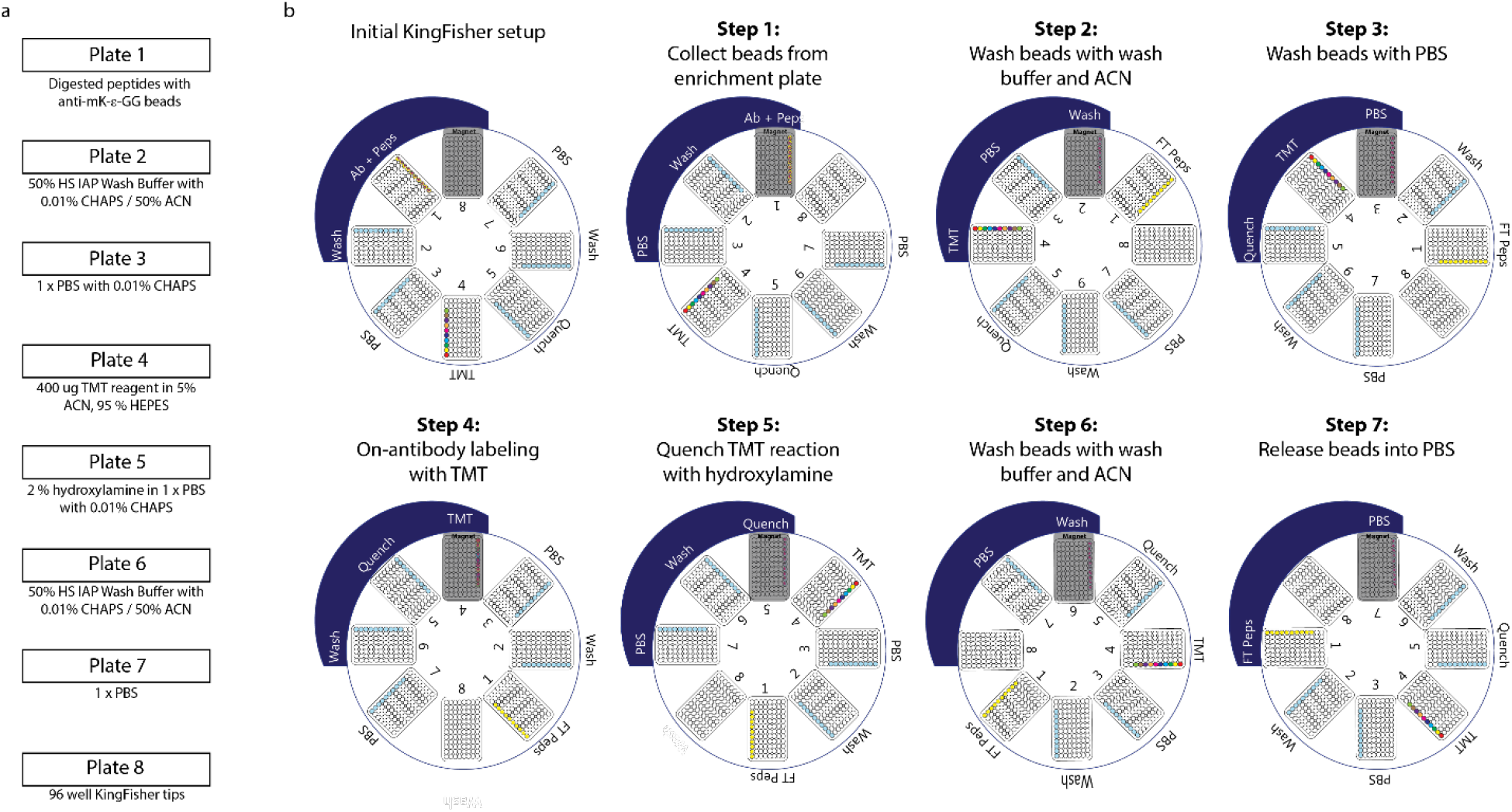
Setup and processing steps for using the KingFisher Flex to automate ubiquitin enrichment. a Plate contents for each of the 8 96-well KF plates. b Overview of each step in the automated ubiquitin enrichment process.

**Supplemental Figure 3.**
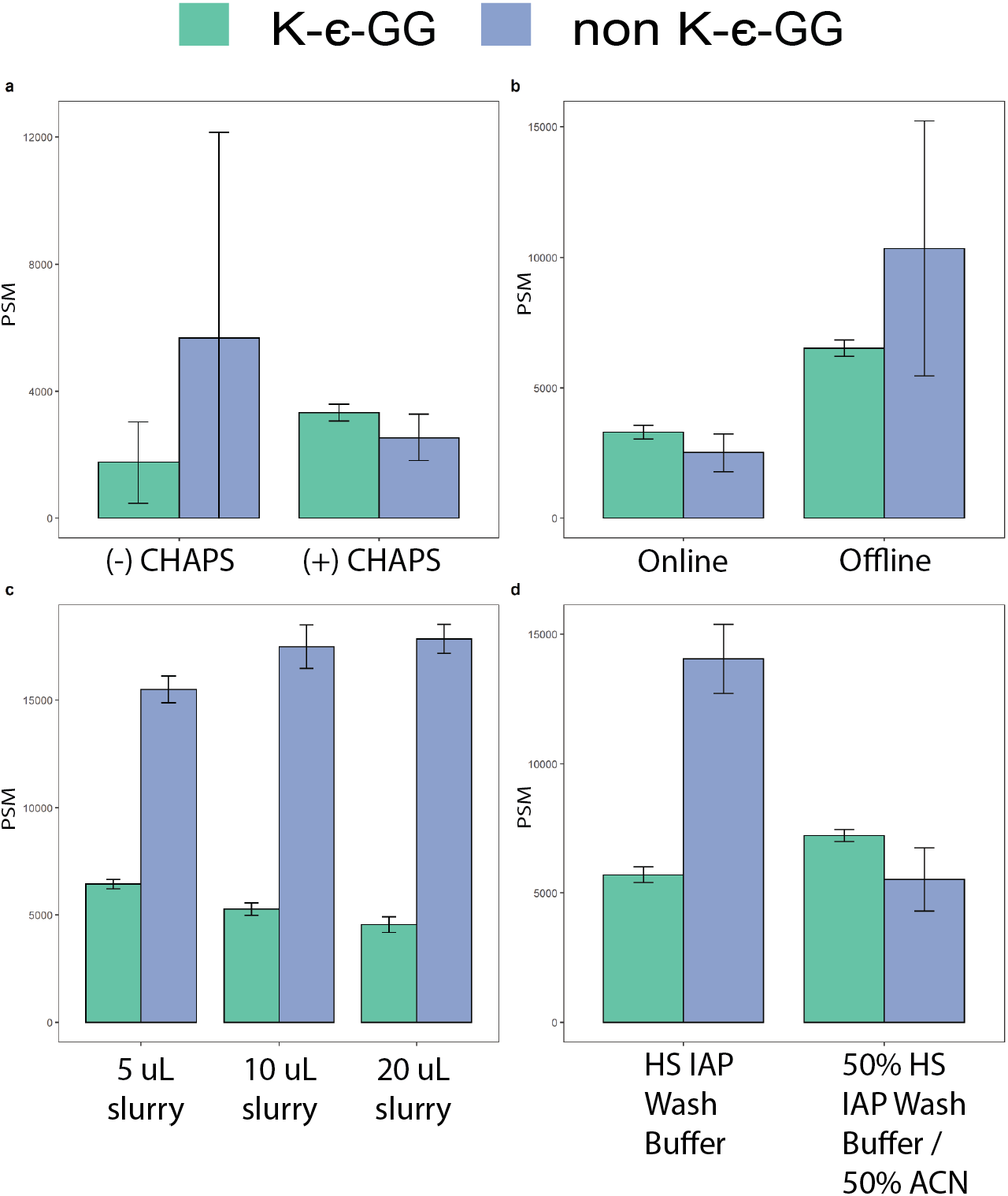
Optimization of bead amounts and washing buffer for K-ε-GG enrichments using the KingFisher Flex instrument. a Bar plots showing identification of K-ε-GG PSMs when 0.01% CHAPS is either absent or present from buffers used on the KingFisher platform. b Bar plots showing identifications of K-ε-GG PSMs from enrichments where the binding of K-ε-GG peptides occurred for 1 hour on the KF robot at RT or 1 hour with end-over-end mixing at 4 °C. **c** Bar plots show identification of K-ε-GG and non K-ε-GG PSM for enrichments using varying amounts of beads (n=5). **d** Bar plots show the identification of K-ε-GG and non K-ε-GG PSM from enrichments with either HS IAP Wash Buffer or HS IAP Wash Buffer diluted 1:1 with ACN.

